# Discovery of A Small Molecule non-IMiD Degrader of ZBTB7A for the Treatment of β-hemoglobinopathies

**DOI:** 10.1101/2025.09.17.676148

**Authors:** Jun Liu, Ziyang Shen, Shin-Young Park, Yanhua Dong, Na Yu, Jing Zeng, Hannah Lee, Braden Pate, Sophia Adamia, Kim Vanuytsel, Junyan Zhang, Shang-Chuen Wu, Adelaide Herman, Shiva Moein, Wenhui Li, Miao Liu, Chong Gao, Xi Tian, Zhaoji Liu, Junsu Kwon, Kunhua Qin, Christoph Budjan, Po-Shen Ko, Claire Shao, Chaitanya Jaladanki, Jiajia Li, Ernie Lee, Bee-hui Liu, Sean Stowell, John P. Manis, David Justus, Gerd A Blobel, Hongbo R Luo, Roger Belizaire, Yi Zheng, Sahand Hormoz, Sarah Nikiforow, Jose A. Cancelas, Hao Fan, Daniel E. Bauer, Daniel Tenen, Li Chai

**Author notes:** Co-first authors (Jun Liu, Ziyang Shen, Shin-Young Park, Yanhua Dong). Co-corresponding **Co-corresponding authors** Jun Liu and Li Chai.

## Abstract

Sickle cell disease and β-thalassemia, two major β-hemoglobinopathies, pose significant clinical challenges globally. Current treatments often face limitations in efficacy and tolerability. The transcription factor ZBTB7A has emerged as a promising therapeutic target for reactivating fetal hemoglobin expression. Here, we report the discovery and characterization of SH6, a small molecule non-IMiD degrader of ZBTB7A. SH6 induces fetal hemoglobin in erythroid cell lines in a CRBN and ZBTB7A-dependent manner, and it is capable of inducing fetal hemoglobin expression in healthy donor, SCD and β-thalassemia patient CD34+ cell derived erythroid cells. The efficacy of SH6 is confirmed in a xenotransplantation humanized mouse model. SH6 outperforms currently available therapeutic agents *in vitro*, and shows synergy with hypomethylating agents. SH6 exhibits a favorable *in vivo* toxicity profile. Our findings establish SH6 as a promising therapeutic lead candidate for further optimization towards clinical development for treatment of sickle cell disease and β-thalassemia.

## Introduction

Sickle cell disease (SCD) and transfusion dependent β-thalassemia are severe inherited hemoglobinopathies caused by mutations in the β-globin gene, leading to defective or insufficient hemoglobin production. They lead to significant clinical morbidity and represent a major health burden worldwide^1^. Current treatments, including γ-globin inducers (e.g. hydroxyurea), transfusions, iron chelation, and bone marrow transplantation, often have limitations in efficacy, tolerability, and/or accessibility ^2^. Novel gene therapies, such as CRISPR/Cas9-mediated disruption of BCL11A, have shown promise but face challenges related to cost, scalability, and long-term safety concerns. As a result, there is an urgent need for more accessible therapeutic strategies that can induce fetal hemoglobin (HbF) as a compensatory mechanism to ameliorate disease severity ^3,4^.

The transcription factor ZBTB7A (also known as leukemia/lymphoma-related factor or LRF) has emerged as a critical repressor of γ-globin expression during the fetal-to-adult hemoglobin switch ^5^. ZBTB7A belongs to the BTB/POZ domain-containing protein family and contains four C2H2-type zinc fingers that mediate its DNA-binding activity. Biochemical and structural studies have shown that ZBTB7A binds directly to the −200 region of the γ-globin (*HBG1* and *HBG2*) promoters via its zinc finger domains, silencing transcription by recruiting corepressors (e.g., NuRD complex) to establish a repressive chromatin state ^6^. Naturally occurring mutations in this binding site are associated with hereditary persistence of fetal hemoglobin (HPFH), underscoring its functional significance in globin regulation ^7^. Genetic perturbation of ZBTB7A or ZBTB7A binding of *HBG1/2* promoters in erythroid cell lines, erythroid cells derived from primary human CD34+ cells, as well as animal models lead to robust reactivation of γ-globin expression without impairing erythroid differentiation ^8,9^. Importantly, ZBTB7A functions independently of BCL11A, another well-characterized γ-globin repressor, making it an attractive alternative therapeutic target for HbF induction.

Transcription factors like ZBTB7A have historically been considered “undruggable” due to their lack of enzymatic activity and intranuclear localization. Recent advances in targeted protein degradation technologies, such as molecular glues, PROTACs, and nanobodies, have opened new avenues for targeting transcription factors like ZBTB7A ^10^. Molecular glue degraders co-opt E3 ubiquitin ligases to recruit specific protein targets for ubiquitination and subsequent proteasomal degradation. This approach has been successfully applied to degrade zinc-finger transcription factors such as IKZF1/3 using immunomodulatory drugs (IMiDs) like lenalidomide ^11^. Building on these advances, molecular glue degraders targeting other repressors of HbF have recently been developed. For example, dWIZ-1 and dWIZ-2 are small-molecule degraders of WIZ (a newly identified HbF transcriptional repressor) based on an IMiD backbone^12^. These studies provide proof-of-concept for using small-molecule degraders to reactivate fetal globin expression and highlight the therapeutic potential of this approach.

In this study, we report the discovery and characterization of SH6, a novel small molecule non-iMID degrader of ZBTB7A. SH6 targets ZBTB7A for CRBN-dependent proteasomal degradation. Using structural insights from the ZBTB7A-DNA complex, we characterized SH6 through *in silico* docking studies. We validated its efficacy in multiple relevant *in vitro* erythroid models and an *in vivo* humanized xenotransplantation mouse model. SH6 demonstrated robust γ-globin induction in HUDEP-2 cells, healthy donor (HD)-derived CD34+ hematopoietic stem and progenitor cells (HSPCs), and patient CD34+ cell-derived erythroid cells from SCD and β-thalassemia patients. Importantly, SH6 induced HbF *in vivo* using a xenotransplantation model. SH6 did not impair erythroid differentiation, cause globin chain imbalance, or exhibit significant toxicity at effective doses tested. SH6 offers a novel therapeutic approach for β-globinopathies with potential advantages over current treatments.

## Methods

### HUDEP-2 cell culture and erythroid differentiation

HUDEP-2 cells were cultured in StemSpan SFEM II media (Stem Cell Technologies) supplemented with 50 ng/mL human stem cell factor, 3 U/mL erythropoietin, 1 μM dexamethasone, and 1 μg/mL doxycycline. For differentiation, cells were cultured in Iscove’s Modified Dulbecco’s Medium (IMDM) supplemented with 5% human AB serum, 3 U/mL erythropoietin, 330 μg/mL holo-transferrin, 10 μg/mL recombinant human insulin, and 2 U/mL heparin. Differentiation protocol is adapted from published protocols ^13^. ZBTB7A−/− and CRBN−/− HUDEP-2 cells were generated using CRISPR-Cas9 ribonucleoprotein (RNP) complexes as previously described ^5,14^(courtesy of Orkin lab).

### Healthy donor and patient-derived CD34 cells

Cryopreserved CD34+ cells from mobilized peripheral blood were obtained from multiple healthy donors (HDs), specifically G-CSF-mobilized, CD34-enriched stem cells sourced from the Fred Hutchinson Cancer Research Center. CD34+ cells from de-identified patients with SCD and β-thalassemia were isolated from whole blood/apheresis specimens provided by the Kraft Family Blood Donor Center and/or Dana Farber Cancer Institute Cell Manufacturing Core Facility in accordance with the Declaration of Helsinki and institutional ethical guidelines (IRB:2025P000309).

### Isolation of CD34 cells from apheresis or peripheral blood specimens

CD34+ hematopoietic stem and progenitor cells were isolated from patient peripheral blood or apheresis products using the EasySep™ Human CD34 Positive Selection Kit (STEMCELL Technologies), following the manufacturer’s instructions. Peripheral blood mononuclear cells (PBMCs) were collected via density gradient centrifugation using Lymphoprep™ (STEMCELL Technologies). The resulting CD34+ cell fraction, typically achieving purity (up to 80%), was immediately available for downstream applications, including flow cytometry, cell culture, or cryopreservation for banking.

### CD34 erythroid differentiation

Healthy donor/patient CD34 cells were thawed and recovered to EDM (IMDM supplemented with 330 μg/mL Holo-Human Transferrin, 10 μg/mL Recombinant Human Insulin, 2 IU/mL Heparin, 5% Inactivated Plasma, 3 lU/mL Erythropoietin, 2 mM L-Glutamine) with three supplements (10^-6 M hydrocortisone, 100 ng/mL SCF, 5 ng/mL IL-3) for 7 days to allow erythroid differentiation, then further differentiated in EDM with one supplement (100 ng/mL SCF) for another 4 days, then further differentiated in EDM with no supplement for 5-7 days. Differentiation protocol is adapted from published protocols ^15^.

### SCD iPSCs

Sickle cell patient-derived iPSCs were obtained from a sickle cell disease-specific iPSC library^16^ and differentiated into erythroid cells using a stepwise suspension protocol as previously described ^17^.

### Western blotting

Cells were lysed in RIPA buffer supplemented with protease inhibitors. Lysates were resolved on 4-12% Bis-Tris gels, transferred to PVDF membranes, and probed with antibodies against ZBTB7A (CST, 50565) and β-actin (Santa Cruz, sc-47778). Blots were developed using Biorad Chemidoc MP Imaging system. Quantification of WB signal performed by ImageJ (v1.54p).

### qRT-PCR

RNA was isolated using the RNeasy Mini Kit (Invitrogen) and reverse transcribed using the High-Capacity cDNA Reverse Transcription Kit (Applied Biosystems). qPCR was performed using TaqMan Gene Expression Assays for *HBG1/2*, HBB, HBA, HBE, and GAPDH on a QuantStudio Real-Time PCR System. qPCR primer sequences are included in Supp Table 1.

### Flow cytometry

Differentiated HUDEP-2 and/or human CD34-derived cells were fixed, permeabilized (BD, 554714), and stained with antibodies against γ-globin (Invitrogen, MHFH01), CD235a (glycophorin A) (BD, 562938), CD71 (Biolegend, 334122), and CD36 (Miltenyi, 130-110-879). Data were acquired on a BD Cantos, CyTek flow cytometer (Cytek Northern Lights™) or Stragedigm S1000Exi cytometer and analyzed using FlowJo (v10).

### HPLC

Globin chains were separated by cation-exchange HPLC on a PolyCAT A column (PolyLC) (Biorad). Samples were eluted with a linear gradient of sodium chloride and sodium phosphate buffers. Peak areas were used to quantify globin chain ratios using the manufacturer’s settings.

### Cellular Thermal Shift Assay (CETSA)

The cellular thermal shift assay (CETSA) was conducted as described by Jafari et al. ^19^ and adapted to our system. For each assay, H1299 cells were seeded at 10^5^ cells/mL in T75 flasks and cultured until ~70% confluency (approximately 3–4 days). Cells were treated with 20 μM SH6 or an equal volume of DMSO (vehicle control) for 2 hours at 37 °C in a humidified incubator with 5% CO_2_. After treatment, cells were pelleted by centrifugation and washed twice with PBS. The cell pellet was resuspended in PBS and aliquoted into equal 100 μL portions. Each aliquot was subjected to a defined temperature (ranging from 44 °C to 68 °C) using a thermal cycler for 3 minutes. After thermal treatment, the samples were lysed in RIPA buffer supplemented with protease inhibitors, followed by sonication. Lysates were clarified by centrifugation at 20,000 × g for 10 minutes at 4 °C to isolate the soluble protein fraction. Supernatants were analyzed by SDS-PAGE and western blotting using antibodies specific for ZBTB7A, CRBN, CUL4A, CUL4B and β actin.

### Human CD34 cell xenotransplantation

NOD.Cg-KitW^−41^J Tyr + Prkdc^scid^Il2rg^tm1^W^jl^/ThomJ (NBSGW, Stock No: 026622) mice (6-8 weeks of age) were purchased from Jackson Laboratory (Bar Harbor, ME; https://www.jax.org) and maintained as homozygotes. Mice were housed in a pathogen-free facility and handled according to the animal protocol, which was approved by the Boston Children’s Hospital’s Animal Care and Use Committee.

Human CD34+ hematopoietic stem and progenitor cells (HSPCs) were transplanted into immunodeficient NBSGW mice, which were conditioned with low-dose busulfan (10 mg/kg). Following confirmation of stable, high-level human engraftment after week 12-16 by flow cytometric analysis of peripheral blood (hCD45+ chimerism) every 4 weeks, mice were randomized to receive vehicle or SH6 treatment (2.5 or 5 mg/kg, intraperitoneally, daily) for three weeks. At the study endpoint, comprehensive analyses were performed on peripheral blood and bone marrow. Two cohorts of mice were pooled for final analysis.

Engraftment, lineage composition, and erythroid differentiation were assessed by flow cytometry using human-specific antibodies. Human erythroid cells (hCD235a+) were isolated from bone marrow by magnetic selection (hCD235a (Glycophorin A) MicroBeads, Miltenyi) for hemoglobin analysis by HPLC and flow cytometry. ZBTB7A protein and globin gene expression analyses were performed on hCD235a-selected bone marrow cells to assess target engagement and globin switching. Peripheral blood was collected for complete blood counts (CBC) and serum chemistry to evaluate hematologic and organ toxicity. Major organs (kidneys, lungs, liver, spleen, and heart) were weighed and sectioned for H&E staining to assess organ-specific toxicity.

### *In silico* docking analysis

Molecular docking analysis was carried out using the Glide module in Schrödinger^20^. The crystal structure of ZBTB7A (PDB ID: 7N5W) was prepared with the Protein Preparation Wizard, and potential ligand-binding sites were identified using the SiteMap program, which evaluates site size, enclosure, solvent exposure, hydrophobicity, hydrogen-bonding capacity, and overall druggability. SH6 was docked into the predicted binding pockets, and key protein-ligand interactions were characterized based on docking scores and binding poses.

To further predict the importance of individual amino acid residues for ligand binding, mutational analysis was performed using the Residue Scanning tool in Schrödinger. Residues within 5.0 Å of the ligand were systematically mutated, and changes in binding affinity (ΔΔG) were calculated using MM-GBSA refinement for both wild-type and mutant complexes. This approach enabled the identification of critical residues that stabilize SH6 binding, providing mechanistic insight into the molecular interaction between SH6 and ZBTB7A.

### Mass Spectrometry

SNU398 cells were treated with SH6 (50 μM) or DMSO for 24 hours and harvested in quadruplicates using RIPA buffer (Sigma) per the manufacturer’s instructions. A total of 200 μg protein from each sample was processed for mass spectrometry as previously described. Samples were boiled at 95 °C and separated on a 12% NuPAGE Bis-Tris precast gel (Thermo Fisher Scientific) at 170 V for 15 minutes in 1× MOPS buffer. Gels were fixed using the Colloidal Blue Staining Kit (Thermo Fisher), and each lane was cut into two equal fractions. In-gel digestion involved destaining in 25 mM ammonium bicarbonate with 50% ethanol, reduction with 10 mM DTT at 56 °C for 1 hour, and alkylation with 55 mM iodoacetamide for 45 minutes in the dark. Proteins were digested overnight at 37 °C in 50 mM ammonium bicarbonate with 2 μg trypsin (Promega). Peptides were desalted using StageTips and analyzed by nanoflow LC-MS/MS on an EASY-nLC 1200 system coupled to a Q Exactive HF mass spectrometer (Thermo Fisher). Separation was done on a 25-cm C18-reversed phase column (75 μm ID) packed in-house with 1.9 μm ReproSil-Pur C18-AQ resin (Dr Maisch), maintained at 40 °C. Peptides were eluted over a 215-minute gradient from 2 to 40% acetonitrile in 0.5% formic acid at 225 nL/min. The Q Exactive HF operated in TOP20 data-dependent acquisition mode, with full MS scans at 60,000 resolution (20 ms max injection time) and MS/MS scans at 15,000 resolution (50 ms max injection time). Raw files were processed with MaxQuant (v1.5.2.8) using label-free quantification (LFQ) on unique plus razor peptides with a minimum of two ratio counts. Carbamidomethylation was set as a fixed modification, while methionine oxidation and N-terminal acetylation were set as variable modifications. Results were filtered at a 1% false discovery rate, and known contaminants, site-only identifications, and reverse hits were excluded. Differential protein analysis was performed using thresholds of log2 fold change < −2 and adjusted p-value < 0.05.

### MTT Cell Viability Assay of SH6-Treated Human Pluripotent Stem Cells

Human embryonic stem cells (hESCs) or human induced pluripotent stem cells (hiPSCs) were seeded into 96-well flat-bottom plates pre-coated with vitronectin-N (VTN-N; Thermo Fisher Scientific, A14700) at 0.5 µg/cm 2, at an initial density of 2,000 cells per well. Cells were cultured in NutriStem hPSC XF medium (Sartorius, 05-100-1B) supplemented with 10 µM Y-27632 dihydrochloride (ROCK inhibitor; Tocris) for the first 24 h after seeding. At 24 h, cells were treated with SH6 dissolved in DMSO to final concentrations of 100 nM, 50 nM, 25 nM, 10 nM, or with vehicle (0.05% v/v DMSO). Culture medium containing SH6 or DMSO was refreshed every 24 h for two additional days (total exposure 72 h). After 72 h of SH6 or vehicle exposure, 10 µL of MTT stock (5 mg/mL in PBS) was added to each well containing 100 µL of medium (final ~0.45 mg/mL MTT), and plates were incubated 3–4 h at 37 °C, 5% CO_2_, protected from light. Formazan crystals were solubilized by carefully removing the medium and adding 100 µL DMSO per well, mixing by gentle pipetting, and incubating for 15 min at 37 °C. Absorbance was recorded at 570 nm with a 630 nm reference, and background from cell-free wells (medium + MTT + DMSO) was subtracted. Viability was expressed as a percentage of the vehicle (0.05% DMSO) control. For each condition, n = 6 wells were measured, and the mean ± SD was reported.

### Statistical analysis

Data are presented as mean ± SD. Minimal two to three biological replicates were included wherever possible, except patient-derived CD34+ cell differentiation, where certain end points were limited to one biological replicate due to limited cellular materials. In the case of qPCR, each biological replicate contains 3 technical replicates. P-values were calculated by two-tailed unpaired t-test using GraphPad Prism (v10.5). *p* < 0.05 was considered statistically significant and marked with *, *p* < 0.01 was marked with **, *p* < 0.001 was marked with ***, *p* < 0.0001 was marked with ****

## Results

### SH6 treatment leads to Decreased ZBTB7A Protein Expression and SH6 Demonstrates Potential Binding to ZBTB7A via *in silico* Docking Analysis

SH6 was previously identified as a non-IMiD SALL4 molecular degrader and is currently under pre-clinical development for treating SALL4-high solid tumors such as Hepatocellular Carcinoma (HCC) ^18^. SH6 was noted to decrease the protein level of other Zinc Finger transcription factors in SNU398 cells via Mass Spectrometry (Fig 1A). ZBTB7A and ZBTB11 were among highly downregulated Zinc Finger transcription factors that contain Zinc Finger domains with substantial homology to the SALL4 Zinc Finger cluster 4 degron by amino acid sequence alignment (Fig S1). SH6 was further evaluated for its ability to modulate ZBTB7A protein levels, given the significant role of ZBTB7A in regulating *HBG1/2* expression and potential as a therapeutic target for treating β-hemoglobinopathies. Mass Spec demonstrated a decrease in relative abundance of ZBTB7A in SH6-treated SNU398 cells (4.1 fold decrease, *p* < 0.01) (Fig 1A’), confirmed by WB analysis (Fig 1B).

**Fig 1.**
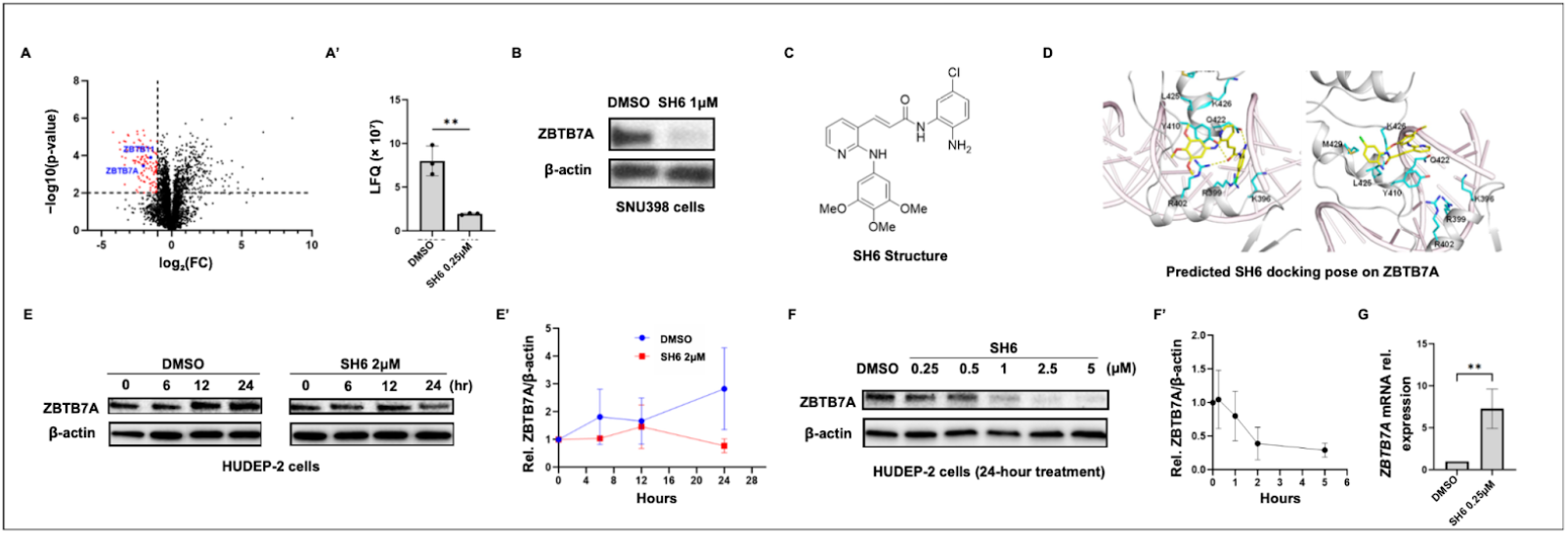
SH6 treatment leads to decreased ZBTB7A protein levels. (A). Mass Spectrometry analysis of SH6 in SNU398 cells identified multiple down-regulated transcription factors, including ZBTB7A and ZBTB11. (B) WB analysis of ZBTB7A protein level in SNU398 cells treated with 1 μM SH6 for 24 hours. (C) Chemical structure of SH6. (D) Predicted docking poses of SH6 against ZBTB7A and predicted essential amino acid residues for Drug-Ligand interaction. (E) Time-dependent effects of ZBTB7A levels (WB) as a response to SH6 treatment in HUDEP-2 cells during differentiation, over 6, 12 or 24 hours (2 μM). E’: Quantification of relative ZBTB7A/Beta Actin signal by FIJI (n=2 biological replicates). (F) Dose-dependent effects of ZBTB7A level (WB) in response to SH6 treatment over 24 hours in HUDEP-2 cells during differentiation. F’ E’: Quantification of relative ZBTB7A/β-Actin signal by FIJI (n=2 biological replicates). (G) qPCR analysis of ZBTB7A mRNA level in response to 0.25 μM SH6 treatment (n=3 biological replicates, each with 3 technical replicates).

Molecular docking and structural analysis provided insights into the interaction between SH6 and ZBTB7A. The known molecular structure of SH6 (Fig 1C) and ZBTB7A protein structure (PDB ID: 7N5W), which includes residues 380-500 encompassing all four C2H2-type zinc fingers, were used for computational modeling. This region is critical for DNA binding, with a preference for the 5’GGGTCA3’ motif, and includes the zinc finger domains essential for transcriptional repression. Binding site prediction using SiteMap, identified two potential binding sites on ZBTB7A (Fig 1D). The first predicted site is located between zinc fingers 1 and 2 (zf1-zf2), while the second is situated between zinc fingers 2 and 3 (zf2-zf3). Both sites exhibited favorable druggability scores, suggesting their suitability for small molecule binding. SH6 was docked into these predicted sites, yielding comparable docking scores of −7.12 and −6.48 for site 1 and site 2, respectively. In the primary binding mode at site 1, SH6 formed key interactions with residues R402, Y410, and Q422, which were identified as critical for ligand stabilization. Notably, the docking poses of SH6 remained consistent in both the presence and absence of DNA, suggesting robust binding under different physiological conditions.

To further investigate these computational predictions, *in silico* mutational analysis was performed on residues within a 5 Å radius of the docked SH6 in both binding sites. Each residue was systematically mutated to all other amino acids, and changes in binding free energy were measured by MM/GBSA calculations.. Mutations at R402, Y410, and Q422 in site 1 significantly destabilized SH6 binding, confirming their importance for ligand interaction. Similarly, mutations at residues in site 2 also affected binding stability but to a lesser extent. Protein stability studies on these mutated residues further supported the computational findings. These results underscore the feasibility of SH6 binding to ZBTB7A and provide a structural basis for its mechanism of action as a possible degrader of ZBTB7A.

We hypothesized that SH6 could similarly reduce ZBTB7A protein levels in erythroid cells. HUDEP-2 cells are an immortalized umbilical cord CD34 cell-derived cell line capable of erythroid differentiation using a well-established three-phase differentiation protocol^21^. They exhibit low baseline γ-globin expression ^21^ and high levels of ZBTB7A expression during the expansion phase, and the early phase of differentiation (EDM2), but with decreased levels later on (EDM3 phase) (Fig S2A). We chose HUDEP-2 cells as a model system for the initial characterization of SH6. Using a short 4-day differentiation protocol (2 day EDM2 and 2 day EDM3, later referred to as abbreviated differentiation protocol), we demonstrated that SH6 treatment reduced ZBTB7A protein levels in HUDEP-2 cells within 24 hours of treatment (Fig 1E, 2 μM SH6 vs DMSO), and the reduction of ZBTB7A protein occurred in a dose-dependent manner (Fig 1F, 0.25 μM to 5 μM SH6 vs DMSO). To investigate whether the observed decrease in ZBTB7A protein was due to transcriptional or post-transcriptional mechanisms, qPCR analysis of ZBTB7A mRNA was performed. SH6 treatment during the abbreviated HUDEP-2 differentiation protocol led to an increase in ZBTB7A mRNA levels (7-fold increase, *p* < 0.01) (Fig 1G), suggesting that the reduction in protein levels was likely not due to decreased transcription but rather post-transcriptional degradation or destabilization mechanisms. The increase in mRNA level could be due to compensatory transcriptional upregulation in response to decreased endogenous protein, a phenomenon well described in the context of protein degraders ^18^.

### SH6 Interacts with ZBTB7A and Mediates ZBTB7A Degradation via the CRL4CRBN Complex

To investigate whether SH6 may function as a small molecule degrader, we tested its ability to induce ZBTB7A degradation in the presence and absence of the proteasome machinery. Molecular glue degraders typically act by recruiting their protein targets to E3 ubiquitin ligases, leading to ubiquitination and subsequent proteasomal degradation ^10^. Previously, SH6 was shown to function in a CRBN/CUL4-dependent manner for SALL4 degradation ^18^. Using CRBN deficient (*CRBN*−/−) HUDEP-2 cells generated via CRISPR-Cas9, we tested the hypothesis that SH6 would fail to degrade ZBTB7A in the absence of CRBN. In wild-type HUDEP-2 cells, SH6 treatment reduced ZBTB7A levels significantly compared to DMSO controls, but this effect was completely abrogated in the CRBN knockout (KO) HUDEP-2 cells (Treatment in EDM2 and EDM3 differentiation, 2 days each, Fig 2A). Notably, the baseline ZBTB7A level is lower in CRBN KO HUDEP-2 cells compared to WT HUDEP-2 cells without drug treatment. Next, we confirmed that SH6-mediated ZBTB7A degradation is dependent on proteasomal activity. Co-treatment of differentiating HUDEP-2 cells with SH6 and the proteasome inhibitor bortezomib blocked SH6-induced ZBTB7A degradation (Fig 2B). These findings suggest that SH6 likely promotes the interaction between ZBTB7A and the CRL4CRBN complex, facilitating the ubiquitination and subsequent degradation of ZBTB7A by the proteasome.

**Fig 2:**
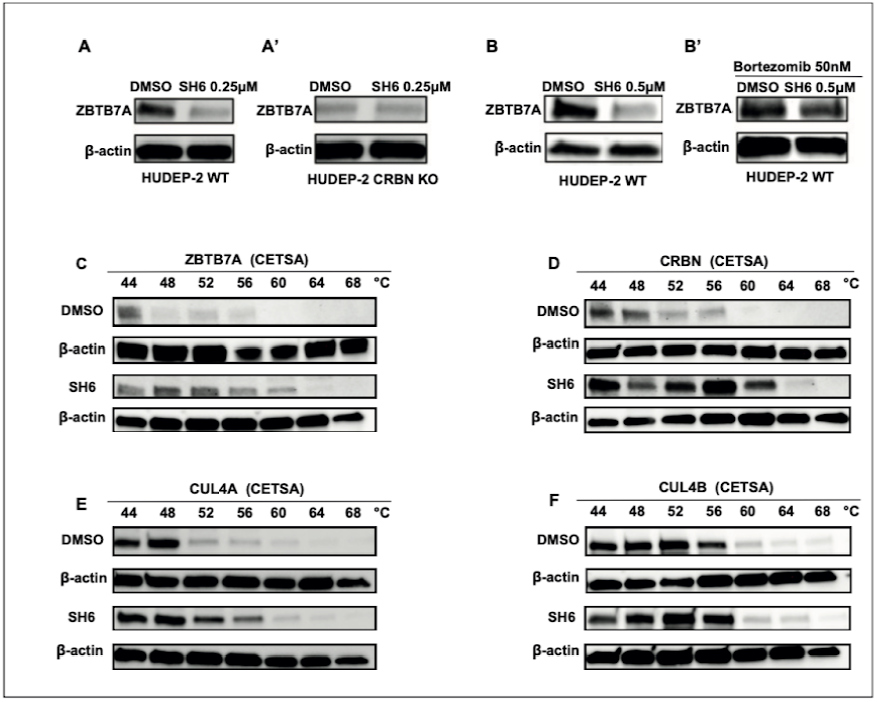
SH6 decreases ZBTB7A in a CRBN/proteasome-dependent manner. (A-A’). WB analysis of ZBTB7A in WT HUDEP-2 cells (A), or CRBN knockout HUDEP-2 cells (A’), treated with 0.25 μM SH6 or DMSO for 24 hours. (B-B’). WB analysis of ZBTB7A in WT HUDEP-2 cells treated with 0.5 μM SH6 or DMSO for 24 hours (B), or in combination with 50 μM Bortezomib (B’). (C) Thermal stability of ZBTB7A in 293T cell lysates pre-treated with DMSO or SH6 at 44, 48, 52, 56, 64, 68 44 °C. (D) Thermal stability of CRBN in 293T cell lysates pre-treated with DMSO or SH6. (E) Thermal stability of CUL4A in 293T cell lysates pre-treated with DMSO or SH6. (F) Thermal stability of CUL4B in 293T cell lysates pre-treated with DMSO or SH6.

To explore the direct interaction between SH6 and ZBTB7A, we employed a cellular thermal shift assay (CETSA). This assay leverages the hypothesis that ligand binding can stabilize proteins against heat-induced denaturation^19^. In untreated cells, ZBTB7A destabilized at 44 degrees Celsius and beyond, but co-treatment with SH6 significantly stabilized ZBTB7A beyond this critical temperature, providing evidence of binding between SH6 and ZBTB7A (Fig 2C). Similar trends are observed with CRBN and CUL4A (i.e. stabilization with SH6 treatment) (Fig 2D,E), but not CUL4B (Fig 2F).

Together, these results establish that SH6 likely acts as a small molecule degrader that binds to CRBN/CUL4A and ZBTB7A, and its mechanism of action likely involves targeting ZBTB7A for proteasomal degradation via the CRL4CRBN complex.

### SH6 Treatment Leads to Significant CRBN/ZBTB7A-dependent Fetal Hemoglobin Induction in HUDEP-2 Cells, and Healthy Donor CD34-derived Erythroid Cells

To evaluate the effects of SH6-mediated ZBTB7A degradation on globin gene expression, we treated HUDEP-2 erythroid cells with SH6 during the abbreviated differentiation protocol. Flow cytometric analysis revealed a dose-dependent increase in HbF induction (Fig S2B), and SH6 at 0.25 μM consistently surpasses or comparable to the effects of other globin inducers such as hydroxyurea (HU), pomalidomide (POM), and decitabine (DAC), all at equivalent or lower doses (Fig 3A, 45.4 % F cells with SH6 (*p* < 0.0001) compared to 28.2 % with HU (*p* < 0.05), 27.2 % with POM (*p* < 0.05), and 39.2 % with DAC (*p* < 0.001)). HbF upregulation was accompanied by a corresponding increase in HbF tetramer frequency, as assessed by HPLC analysis: SH6 treatment increased the HbF% from approximately 3% to 19% of total hemoglobin (*p* < 0.001) (Fig 3B). To ensure the validity of the phenotypes observed in the abbreviated HUDEP-2 differentiation protocol, we followed the conventional 11 day differentiation protocol, and observed robust HbF induction at day 11 with minimal changes in erythroid differentiation markers in HUDEP-2 cells treated with 0.25 μM SH6 (Fig S3).

**Fig 3.**
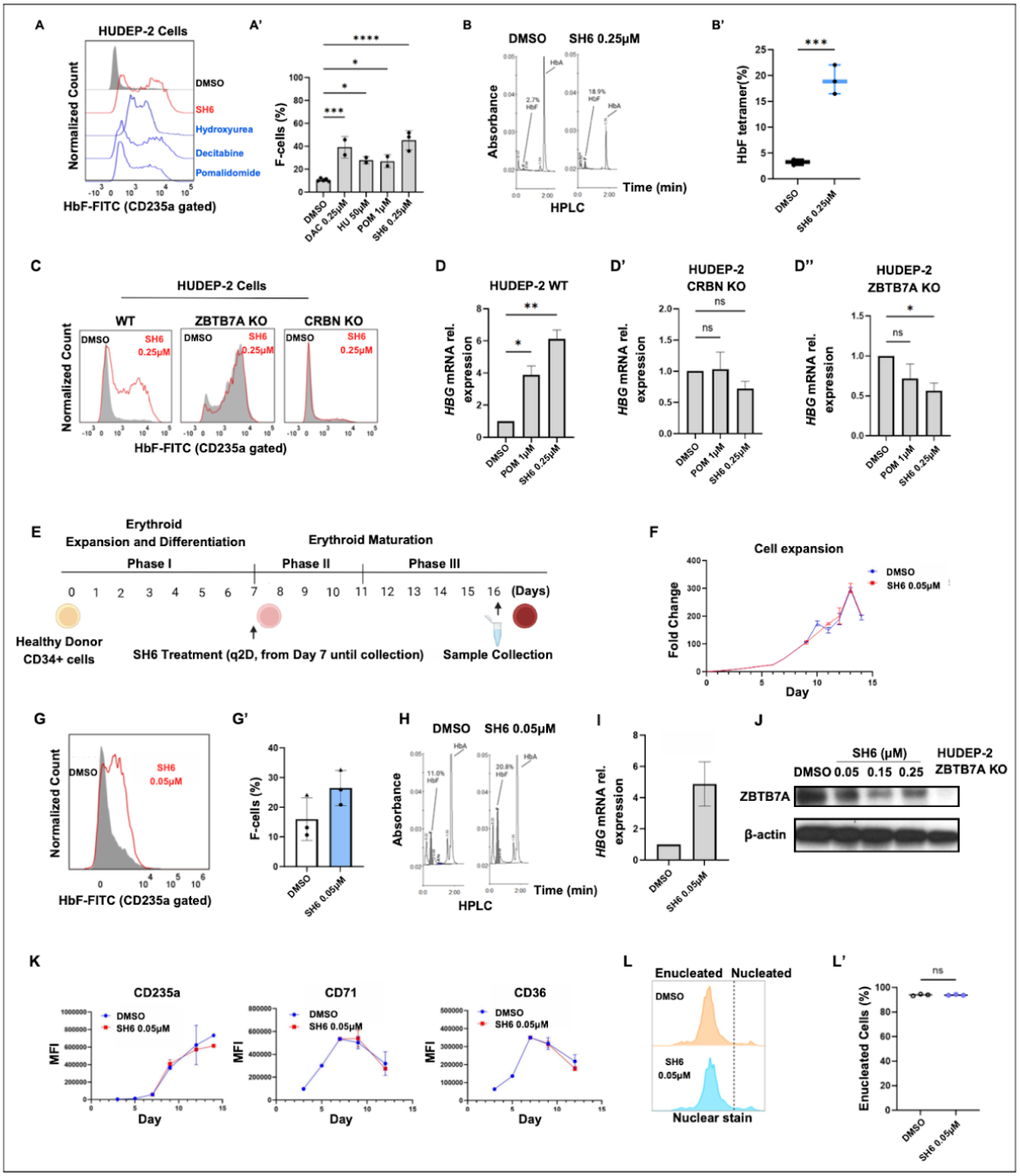
SH6 Leads to CRBN/ZBTB7A-Dependent Fetal Hemoglobin Induction in HUDEP-2 and Healthy Donor CD34-derived Erythroid Cells. (A) Representative flow cytometric analysis of HbF (CD235a+) in HUDEP-2 cells at day 4, with an abbreviated differentiation protocol (2D EDM2 and 2D EDM3), treated with DMSO, Decitabine (DAC), Pomalidomide (POM), Hydroxyurea (HU), and SH6. (A’) Quantification of HbF induction (%F cells among CD235a+ cells) for all treated conditions (n=2 biological replicates). (B) Representative HPLC analysis of HbF in HUDEP-2 cells treated with 0.25 μM SH6 vs DMSO, at day 7 of differentiation. (B’) Quantification of HbF% tetramer of HPLC analysis of HbF in HUDEP-2 cells treated with 0.25 μM SH6 vs DMSO, at day 7 of differentiation. (C) Representative flow cytometric analysis of HbF (CD235a+) in WT, CRBN KO and ZBTB7A KO HUDEP-2 cells at day 4 differentiation, treated with 0.25 μM SH6 vs DMSO. (D) qPCR analysis of relative HBG mRNA levels in WT, CRBN KO and ZBTB7A KO HUDEP-2 cells at day 4 differentiation, treated with 0.25 μM SH6, 1 μM Pomalidomide (positive control) vs DMSO (n=3 biological replicates, each with 3 technical replicates). (E) Schematic of healthy donor CD34+ cell differentiation, drug treatment and sample collection/analysis. (F) CD34+ cell erythroid differentiation cell expansion curve for cells treated with 0.05 μM SH6 or DMSO. (G) Representative flow cytometric analysis of HbF (CD235a+) in differentiated CD34+ cells at day 13 treated with 0.05 μM SH6 vs DMSO (G) Quantification of flow cytometric analysis of HbF (CD235a+) in differentiated CD34+ cells at day 13-15 (n= 2 donors, each with 1-2 biological replicates). (H) Representative HPLC analysis of HbF in differentiated CD34+ cells treated with 0.05 μM SH6 vs DMSO, at day 15 of differentiation. (I) Quantification qPCR analysis of *HBG1/2* mRNA in differentiated CD34+ cells treated with 0.05 μM SH6 vs DMSO (n=1 donor and 1 biological replicate, 3 technical replicates). (J) WB analysis of ZBTB7A protein in differentiated CD34+ cells treated with SH6 vs DMSO (ZBTB7A KO HUDEP-2 lysate = negative control). (K) Quantification of flow cytometric analysis of erythroid differentiation markers (CD235a, CD71 and CD36 MFI) during CD34+ erythroid differentiation, in cells treated with 0.05 μM SH6 vs DMSO (n=1 donor and 3 biological replicates). (L) Quantification of flow cytometric analysis erythroid enucleation (% enucleated cells) during CD34+ differentiation, in cells treated with 0.05 μM SH6 vs DMSO (n=1 donor and 3 biological replicate).

Consistent with the requirement for CRBN in SH6-mediated ZBTB7A degradation, SH6 failed to induce HbF protein in CRBN−/− HUDEP-2 cells (Fig 3C). Similarly, ZBTB7A−/− HUDEP-2 cells, showed no further increase in γ-globin protein upon SH6 treatment, supporting the hypothesis that SH6’s effects depend on intact ZBTB7A protein (Fig 3C). qPCR analysis demonstrates *HBG1/2* upregulation on a transcriptional level in WT HUDEP-2 cells (6.1-fold increase compared to DMSO, more pronounced than *HBG1/2*-inducing effect of POM (*p* < 0.01)), but not in CRBN or ZBTB7A KO HUDEP-2 cells (Fig 3D). Interestingly, CRBN−/− HUDEP-2 cells retained the ability to upregulate HbF when treated with DAC, consistent with the notion that SH6 and DAC act through distinct mechanisms (Fig S4). Notably, co-treatment with DAC and SH6 resulted in additive HbF induction, as demonstrated by HPLC and qPCR (Fig S4). These findings position SH6 as a promising alternative or adjunct to other pharmacologic therapies for β-hemoglobinopathy.

To further assess the therapeutic potential of SH6, we evaluated its effects on γ-globin expression and erythroid differentiation in primary human CD34+ hematopoietic stem and progenitor cells (HSPCs) derived from two healthy donors. CD34+ cells were differentiated using a standard three-phase protocol (Fig 3E), with SH6 treatment initiated on day 7 and continued every other day until harvest between days 14 and 17. SH6 was effective at a nanomolar dose range, with a comparable expansion curve to DMSO-treated cells during differentiation (Fig 3F). 0.05 μM SH6 treatment led to a trend of increase in HbF levels as measured by flow cytometry (Fig 3G) and HPLC (Fig 3H, and Fig S5), and *HBG1/2* mRNA by qPCR (Fig 3I). WB analysis showed ZBTB7A protein levels decrease in a dose-dependent manner (Fig 3J). Markers of erythroid differentiation (CD235a, CD36, and CD71) followed normal trajectories (Fig 3K). Additionally, enucleation rates (93.9% with SH6 compared to 93.9% with DMSO) remained consistent with DMSO-treated controls (Fig 3L).

### SH6 Treatment Leads to Significant Fetal Hemoglobin Induction Sickle Cell and β-Thalassemia Patient-Derived Primary CD34+/iPSC Differentiated Erythroid Cells

To further evaluate the therapeutic potential of SH6 in disease-relevant models, we tested its ability to induce fetal hemoglobin (HbF) in erythroid cells differentiated from primary CD34+ hematopoietic stem and progenitor cells (HSPCs) obtained from sickle cell disease (2 patients) and β-thalassemia patients (2 patients) (Fig 4A). CD34+ cells were isolated from mobilized peripheral blood and differentiated into erythroid cells for analysis. In SCD patient-derived erythroid cells, SH6 treatment at 0.05 μM resulted in a significant increase in γ-globin expression, as demonstrated by flow cytometry (Fig 4B 69% F cells with SH6 vs 49% with DMSO (*p* < 0.01)), qPCR (Fig 4C, 2.3-fold increase in relative HBG mRNA level (*p* < 0.05) compared to DMSO, 50% decrease in relative *HBB/HBB*^*S*^ mRNA level compared to DMSO (*p* < 0.01)), and HPLC (Fig 4D 15.0% HbF with SH6 vs 9.5% HbF with DMSO (*p* < 0.05)). These findings align with observations in healthy donor-derived CD34+ cells, further supporting the robustness of SH6-induced HbF induction. qPCR analysis was consistent with the β-globin switch (increased *HBG1/2* transcription and concurrent decreased *HBB/HBB*^*S*^ transcription). A brief dose-escalating experiment (0.01 μM, 0.05 μM, 0.1 μM) was performed with one single SCD patient’s CD34 cells (Fig 4E, unable to perform with the second patient due to scarcity of cells). A dose-dependent HbF induction was observed, and an effect can be observed at as low as 0.01 μM dose (Fig 4E). HPLC showed 9.3%, 11.5%, and 11.8% HbF with 0.01 μM, 0.05 μM, 0.1 μM doses, compared to 8.1% HbF with DMSO (all *p* < 0.0001) (Fig 4E). The trend of HbF elevation persists at terminal differentiation at D19 (Fig S5). While the higher doses appeared to dampen cell expansion curves (Fig 4G), none of the doses tested affected enucleation rates (Fig 4F) or erythroid differentiation markers expression (Fig 4H).

**Fig 4.**
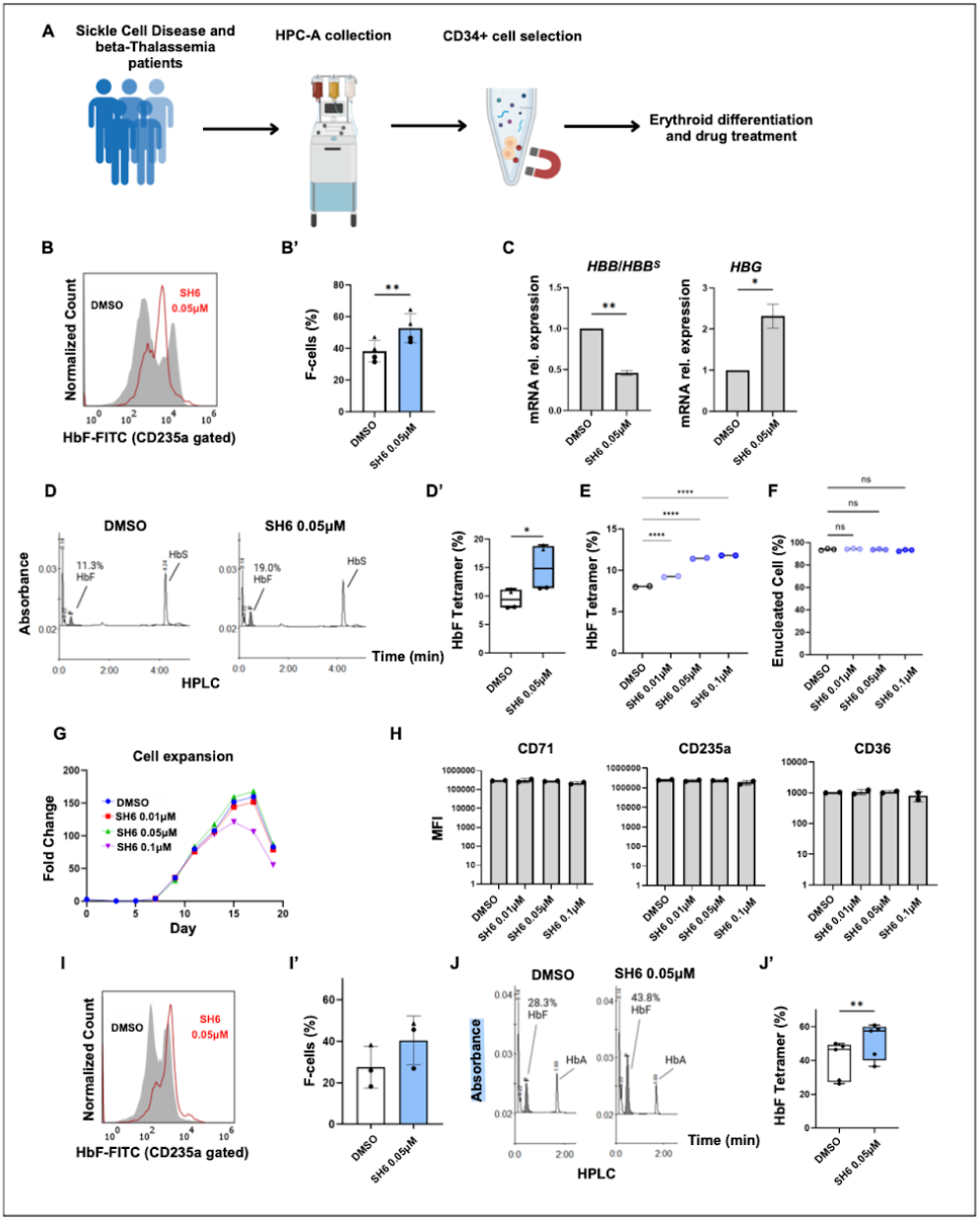
SH6 Induces Fetal Hemoglobin in Sickle Cell Disease and β-Thalassemia Patient-Derived CD34+ Differentiated Erythroid Cells. (A) Schematic of the collection and isolation of CD34+ cells from mobilized SCD and β-halassemia patients. (B) Representative flow cytometric analysis of HbF (CD235a+) in differentiated SCD CD34+ cells at day 15 treated with 0.05 μM SH6 vs DMSO. (B’) Quantification of flow cytometric analysis of HbF (CD235a+) in differentiated SCD CD34+ cells at day 14-16 (n= 2 donors, each with 1-2 biological replicates). (C) Quantification qPCR analysis of *HBG1/2* and *HBB* mRNA in differentiated SCD CD34+ cells treated with 0.05 μM SH6 vs DMSO (n=1 donor and 2 biological replicate, each with 3 technical replicates). (D) Representative HPLC analysis of HbF in differentiated SCD CD34+ cells treated with 0.05 μM SH6 vs DMSO, at day 15 of differentiation. (D’) Quantification HPLC analysis of HbF in differentiated SCD CD34+ cells treated with 0.05 μM SH6 vs DMSO, day 15-17 of differentiation (n= 2 donors, each with 1-2 biological replicates). (E) An abbreviated dose response effect of HbF (quantified by HPLC) at D17 of differentiation, from a single SCD patient’s CD34+ differentiation (n=1 donor, with 2 biological replicates). (F) An abbreviated dose response quantification of erythroid enucleation (% enucleated cells, flow cytometric analysis) from a single SCD patient’s CD34+ differentiation at D19 (n=1 donor, with 3 biological replicates). (G) CD34+ cell erythroid differentiation cell expansion curve for cells treated in the abbreviated dose-response experiment as described in E and F. (H) Quantification of flow cytometric analysis of erythroid differentiation markers (CD235a, CD71 and CD36 MFI) for cells treated in the abbreviated dose-response experiment as described in E and F, at Day 17. (I) Representative flow cytometric analysis of HbF (CD235a+) in differentiated β-Thal CD34+ cells at day 15 treated with 0.05 μM SH6 vs DMSO. (I’) Quantification of flow cytometric analysis of HbF (CD235a+) in differentiated SCD CD34+ cells at day 14-16 (n= 2 donors, each with 1-2 biological replicates). (J) Representative HPLC analysis of HbF in differentiated β-Thal CD34+ cells treated with 0.05 μM SH6 vs DMSO, at day 15 of differentiation. (J’) Quantification HPLC analysis of HbF in differentiated β-Thal CD34+ cells treated with 0.05 μM SH6 vs DMSO, day 15-17 of differentiation (n= 2 donors, each with 1-2 biological replicates).

Similarly, in β-thalassemia CD34+ differentiated erythroid cells, SH6 treatment at 0.05 μM increased γ-globin expression. Flow cytometry showed a trend of increase in HbF-positive cells (Fig 4I, 68.4% F cells with SH6, compared to 36.8% F cells with DMSO), and HPLC analysis demonstrated an increase in HbF levels from 28.3% to 43.8% (*p* < 0.01) (Fig 4J), of total hemoglobin with 0.05 μM SH6 treatment. A mini-dose-escalation treatment (0.05 μM and 0.1 μM) was performed with one β-thalassemia patient’s specimen, and similar to SCD, we observed a trend of dose-dependent HbF induction both with flow cytometry and HPLC (data not shown, single biological replicate due to limited specimen).

To further evaluate the clinical relevance of SH6, we sought to test its effects in additional patient genetic backgrounds by leveraging erythroid cells derived from sickle cell patient-specific induced pluripotent stem cells (iPSCs). SH6 treatment increased γ-globin in iPSC-derived erythroid cells from a healthy donor and a SCD patient known to be non-responsive to hydroxyurea (Fig S6).

### SH6 treatment leads to *in vivo* HbF induction with minimal toxicity in xenotransplanted NSGW mice

Healthy donor human CD34+ hematopoietic stem and progenitor cells (HSPCs) were transplanted into two separate cohorts of immunodeficient NBSGW mice (with C-KIT mutation to facilitate human erythroid engraftment), either without conditioning or following low-dose busulfan treatment to enhance engraftment.^22^(Fig 5A). Cohort 1 received transplants of pooled donor cells derived from three individual donors, whereas cohort 2 received transplants of cells from a single donor per mouse, utilizing three donors distinct from those in cohort 1. Engraftment and hematopoietic reconstitution were monitored longitudinally by flow cytometric analysis of peripheral blood, via human (hCD45+) versus mouse (mCD45+) leukocyte chimerism (Fig S7). Engraftment monitoring was performed every four weeks post-transplantation, beginning at week 4 and continuing through at least week 16. Two cohorts of mice were transplanted one month apart, and randomization to three treatment groups (vehicle, 2.5 mg/kg, 5 mg/kg, by daily IP injection for 3 weeks) was performed independently for each cohort (Fig 5A). Treatment started between weeks 16-18. By week 16, robust human hematopoietic engraftment was achieved in the majority of mice with comparable engraftment rates between the two cohorts: the mean percentage of hCD45+ cells in peripheral blood at week 16 was 62.7 for cohort 1, and 69.5 for cohort 2 (Fig S7). Data from two cohorts were combined for analysis (vehicle, 4 mice, 2.5 mg/kg, 6 mice, 5 mg/kg, 5 mice, by daily IP injection for 3 weeks). One mouse was excluded from analysis due to premature death and having received drug treatment for only two weeks.

**Fig 5.**
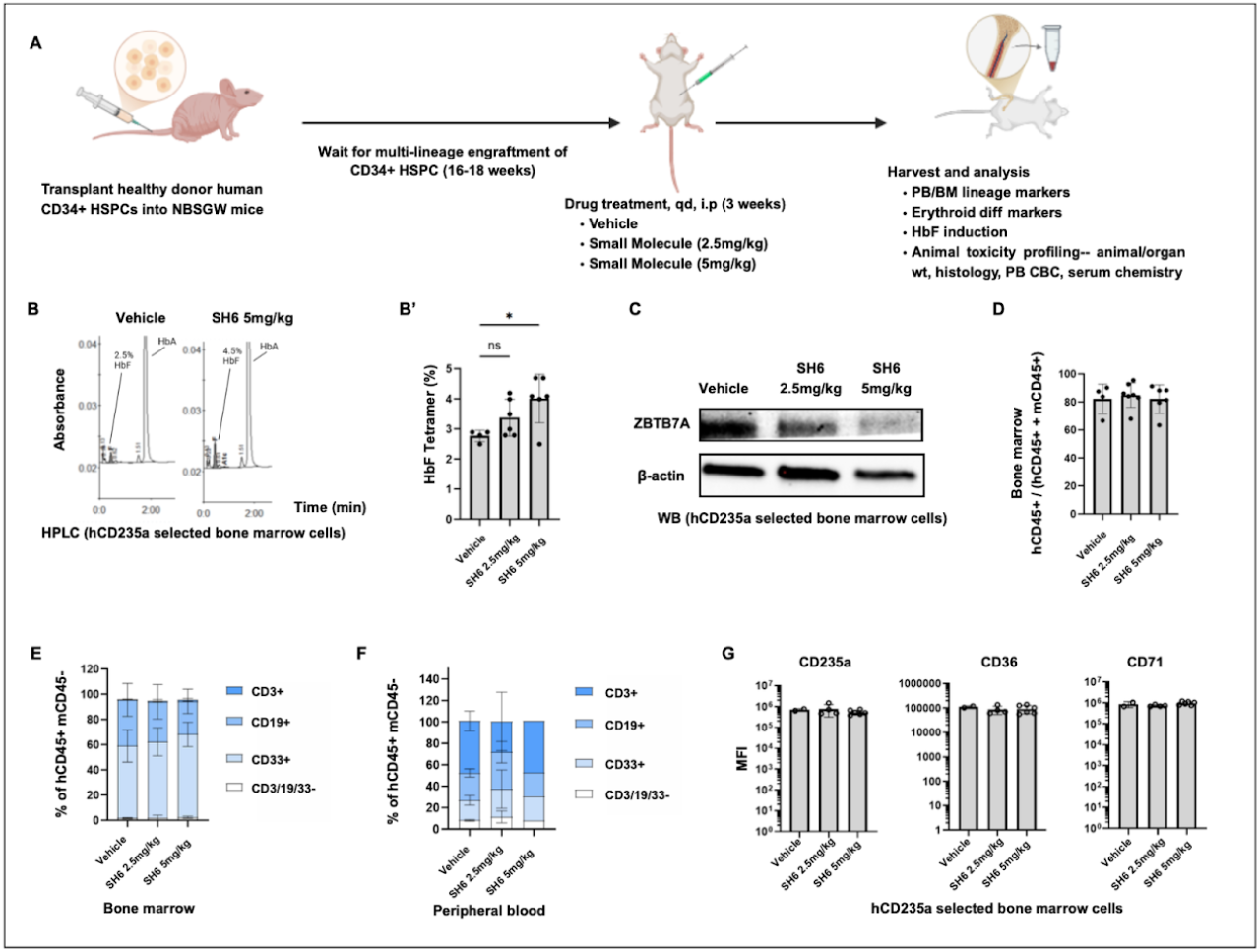
SH6 treatment leads to *in vivo* HbF induction with minimal toxicity in xenotransplanted NBSGW mice. (A) Schematic of murine xenotransplantation of healthy donor CD34+ cells, *in vivo* drug treatment, and specimen harvest/analysis. (B) Representative HPLC analysis of HbF in hCD235a-selected erythroid cells from mouse bone marrow (vehicle vs SH6 5 mg/kg, treated daily for 3 weeks). (B’) Quantification HPLC analysis of HbF in hCD235a-selected erythroid cells from mouse bone marrow (vehicle vs SH6 5 mg/kg, treated daily for 3 weeks) (n= 4 vehicle control, n=6 2.5 mg/kg dose, n=5 5 mg/kg dose). (C) WB analysis of ZBTB7A protein in hCD235a-selected erythroid cells from mouse bone marrow, treated with vehicle, 2.5 mg/kg dose, and 5 mg/kg dose. (D) Bone marrow human CD45+ engraftment across three treatment groups. (E) Whole bone marrow human lymphoid and myeloid lineage marker flow cytometric analysis of xenotransplanted mice, across three treatment groups. (F) Peripheral blood human lymphoid and myeloid lineage marker flow cytometric analysis of xenotransplanted mice, across three treatment groups. Only 1 mouse in the 5 mg/kg group had sufficient peripheral blood from cardiac bleed for lineage marker analysis, after taking specimen for CBC and serum chemistry. (G) Quantification of flow cytometric analysis of erythroid differentiation markers (CD235a, CD71 and CD36 MFI) in hCD235a-selected erythroid cells from mouse bone marrow across three treatment groups.

HPLC analysis of engrafted human erythroid cells (hCD235a-selected bone marrow cells, Fig S8) revealed a small yet significant induction of HbF (Fig 5B, 4.0% HbF with 5mg/kg dose, 2.8% HbF with vehicle (p < 0.05)), suggesting modulation of HbF expression *in vivo*. The trend of HbF induction is also demonstrated by flow cytometric analysis (Fig S8). SH6 exposure resulted in dose-dependent target engagement as demonstrated by reduced ZBTB7A protein levels in human CD235a+ selected bone marrow cells, with greater ZBTB7A protein reduction observed at the 5 mg/kg dose compared to 2.5 mg/kg treatment (Fig 5C).

Bone marrow hCD45 engraftment were comparable across all treatment groups prior to harvest (Fig 5D). Bone marrow and peripheral blood lineage analysis demonstrated multilineage human hematopoietic reconstitution, with comparable percentages of T cells, B cells, and myeloid cells between vehicle and treated groups (Fig 5E,F)^23^. Flow cytometric immunophenotyping of erythroid differentiation markers in hCD235a+ selected bone marrow cells showed consistent expression of key surface markers CD235a, CD36, and CD71 across all treatment groups, indicating preservation of normal erythroid maturation programs *in vivo* despite target modulation (Fig 5G).

Treatment with SH6 was well tolerated at 2.5 mg/kg and 5 mg/kg as assessed by stable body weight trajectories in both cohorts compared with vehicle control (Fig S7). Comprehensive safety evaluation revealed no significant treatment-related adverse effects on key physiological parameters. Serum chemistry analysis showed normal levels of urea nitrogen, creatinine, albumin, and liver enzymes (AST, ALT) across all treatment groups (Fig S9). Complete blood count parameters, including white blood cell count, red blood cell count, hemoglobin, platelet count, and mean corpuscular volume, remained within comparable ranges across groups (Fig S9). Organ weight analysis of kidneys, lungs, liver, spleen, and heart demonstrated no significant changes compared to vehicle controls, indicating the absence of overt target organ toxicity (Fig S10). Tissue histology with H&E staining also showed normal gross tissue architecture (data not shown). To evaluate the potential cytotoxicity of SH6, we performed viability assays in HD-derived mononuclear cells (MNCs) from three donors (Fig S13, up to 10 μM), and assessed embryotoxicity using both primary and iPS-derived human embryonic stem cells (Fig S14, up to 0.1 μM), both showing favorable toxicity profiles in the dose ranges tested.

These findings establish that the SH6 achieves dose-dependent *in vivo* HbF induction through the reduction of ZBTB7A protein, while maintaining normal hematopoietic function and demonstrating a favorable safety profile in this humanized mouse xenotransplantation model at the treated doses.

## Discussion

In this study, we report the discovery and characterization of SH6, a small molecule non-IMiD degrader of ZBTB7A, a transcriptional repressor of fetal globin. SH6 induces fetal (γ-globin) expression by promoting CRBN-dependent proteasomal degradation of ZBTB7A. It effectively induces γ-globin in healthy donor-derived CD34+ erythroid progenitors, and patient-derived erythroid cells from SCD and β-thalassemia patients, as well as in iPSC-derived erythroid cells. Herein we demonstrated robust γ-globin induction at nanomolar doses via SH6 treatment without impairing erythroid differentiation or causing globin chain imbalance. Our *in vivo* studies showed that SH6 induces HbFexpression in engrafted human erythroid cells, while preserving normal erythroid differentiation and multilineage hematopoiesis, with minimal hematopoietic or organ toxicity. These findings position SH6 as a promising therapeutic candidate for further optimization and clinical development. The synergy observed between SH6 and DAC suggests that combining epigenetic modulators with transcription factor degraders could be a powerful strategy for HbF reactivation therapies.

The CRBN-dependent activity of SH6 is reminiscent of immunomodulatory drugs (IMiDs) such as thalidomide and lenalidomide, which bind CRBN and redirect its substrate specificity^10^. Molecular docking studies revealed that SH6 may bind to a pocket between the zinc finger domains of ZBTB7A, promoting its interaction with CRBN through a novel structural mechanism. This unique binding mode highlights the potential of targeting ZBTB7A with small molecule degraders without the burden of adverse effects associated with other CRBN modulators like IMiDs. Further structural studies are needed to elucidate the precise binding interactions between SH6, ZBTB7A, and CRBN, which could inform the rational design of next-generation ZBTB7A degraders with improved pharmacological properties.

The development of SH6 builds on recent progress in targeted protein degradation. The availability of multiple molecular glue degraders in clinical development highlights the growing interest in transcription factor degradation as a therapeutic strategy for β-globinopathies. Recent advances in molecular glue degraders targeting transcriptional repressors of fetal globin provide additional context for our findings and support clinical feasibility of this therapeutic approach. A newly reported WIZ degrader (Novartis) targets this newly identified fetal globin repressor to induce HbF expression. Like SH6, WIZ degrader acts via CRBN-dependent proteasomal degradation but targets a different regulatory pathway for HbF induction ^12^. Additionally, Bristol Myers Squibb (BMS) has presented data on a dual degrader targeting both ZBTB7A and WIZ that has already entered clinical trials for SCD^25^. Novartis WIZ degrader and BMS dual degrader de-risk our approach of ZBTB7A degradation, by demonstrating clinical feasibility while differentiating our compound based on its specificity and distinct mechanism of action.

### Clinical relevance of SH6 in β-hemoglobinopathy *in vitro* and *in vivo* models

We demonstrated the therapeutic potential of SH6 across clinically relevant models, including primary CD34+ HSPCs from healthy donors, SCD and β-thalassemia patients and iPSC-derived erythroid cells from SCD patients. In both disease contexts, SH6 induced γ-globin expression while reducing β-globin levels, consistent with the expected globin switch mechanism upon ZBTB7A depletion. The iPSC data highlights the potential of SH6 to offer therapeutic benefits to patients non-responsive to hydroxyurea, the first-line, and only FDA-approved HbF-inducing small-molecule drug. Importantly, α-globin levels were unaffected by SH6 treatment, minimizing concerns about globin chain imbalance. In addition to fetal globin induction, SH6 also induces *HBE* (embryonic globin) (Fig S11). Notably, SH6 was effective in an hydroxyurea non-responder iPSC model, suggesting its potential to overcome limitations of standard SCD treatments. This finding is significant given the heterogeneity in patient responses to hydroxyurea and underscores the need for alternative therapeutic approaches. The *in vivo* relevance was confirmed in humanized NBSGW mice, where low doses of SH6 (5 mg/kg/day) induced dose-dependent HbF in bone marrow-derived human erythroid cells with concomitant ZBTB7A degradation. There may be room for further dose-escalation, since SH6 is well tolerated at up to 50 mg/kg dose in a HCC xenotransplantation model ^18^.

### SH6 has favourable toxicity profile *in vitro* and *in vivo*

While our findings highlight the therapeutic potential of ZBTB7A degradation for HbF induction, it is important to consider possible side effects given ZBTB7A’s known roles during and beyond erythropoiesis. ZBTB7A has been implicated in lymphoid lineage specification and myeloid differentiation ^26^. Loss of ZBTB7A can alter lymphoid homeostasis and promote myeloid skewing in hematopoietic stem cells (HSCs) ^26^. Furthermore, ZBTB7A plays roles in other tissues, including tumor suppression in certain cancers ^26^. These functions suggest that systemic degradation of ZBTB7A could have unintended consequences on immune function or tumorigenesis. We observed minimal toxicity across multiple systems *in vitro* and *in vivo* at the doses treated. In the *in vivo* setting, mice body weights remained stable, serum chemistry (creatinine, AST/ALT) showed no significant organ impairment, and organ histology was unremarkable. Complete blood counts indicated only a mild platelet trend reduction. This aligns with *in vitro* observations of maintained erythroid differentiation and enucleation, supporting SH6’s therapeutic window.

In previously published studies using liver cancer mouse xenotransplantation models, SH6 was well-tolerated at doses up to 50 mg/kg/day without significant changes in body weight or organ histology ^18^. Pharmacokinetic studies revealed good oral bioavailability (F = 35%), moderate plasma half-life (t1/2 = 3.5 h), and low clearance (CL = 8 mL/min/kg), supporting its favorable drug-like properties ^18^. Additional preclinical studies are required to evaluate long-term safety and efficacy in mice and non-human primates before advancing toward clinical trials.

### Limitations and Future Directions

While our study establishes the therapeutic potential of SH6 as a ZBTB7A degrader for HbF induction, several questions remain unanswered. First, the precise structural basis for SH6-mediated recruitment of ZBTB7A to CRBN requires further investigation using techniques such as cryo-electron microscopy or X-ray crystallography. Second, although our data suggest minimal toxicity at lower doses *in vitro* models, comprehensive toxicology studies are needed to assess off-target effects in tissues where ZBTB7A plays critical roles. Additionally, while we demonstrated synergy between SH6 and DAC *in vitro*, it will be important to evaluate combination therapies *in vivo* using xenograft models or patient-derived organoids. Finally, expanding preclinical studies across diverse patient backgrounds will help assess variability in response to SH6 treatment.

## Conclusion

In conclusion, this study identifies SH6 as a small molecule degrader targeting ZBTB7A for HbF induction. By degrading ZBTB7A via the CRL4CRBN complex, SH6 achieves γ-globin induction without significantly impairing erythroid differentiation or causing toxicities. Its efficacy across diverse disease models highlights its potential as a therapeutic candidate for β-globinopathies. SH6 serves as a proof of principle that ZBTB7A degraders can be a promising therapeutic approach for β-hemoglobinopathies.

## Acknowledgement

Stuart Orkin lab, Robert Hubbard, Annalisa De Ruscio, Lucrezia Rinaldi, Dana Farber CMCF

## Author Information

Jun Liu, Ziyang Shen, Shin-Young Park and Yanhua Dong contributed equally to this work.

## Co-senior authors

Daniel E. Bauer, Daniel Tenen or Li Chai

