## Supplemental Figures for "Discovery of A Small Molecule non-IMiD Degrader of ZBTB7A for the Treatment of β-hemoglobinopathies"

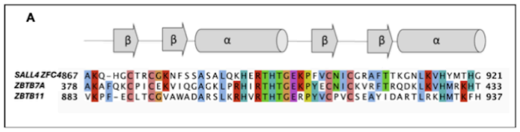


Fig S1. Alignment of the amino acid sequence of published SH6 degron on SALL4 (Zinc Finger Cluster 4) and Zinc Finger domains of ZBTB7A and ZBTB11.

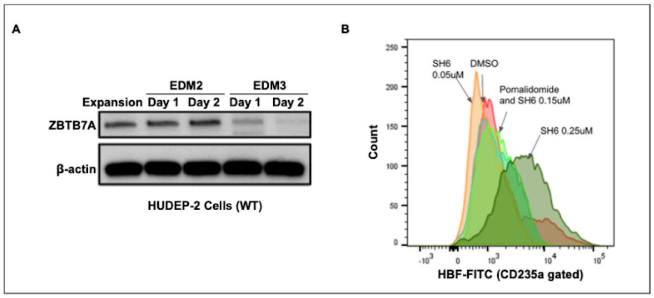


Fig S2. ZBTB7A expression during HUDEP-2 differentiation and dose-dependent HbF induction in WT HUDEP-2 cells. (A) WB analysis of endogenous ZBTB7A protein level during expansion phase and during 2 days of EDM2 and 2 days of EDM3 differentiation. (B) Flow cytometric analysis of dose-dependent HbF induction in WT HUDEP-2 cells (doses treated: SH6 0.05 μM, 0.15 μM and 0.25 μM vs DMSO). POM: Pomalidomide.

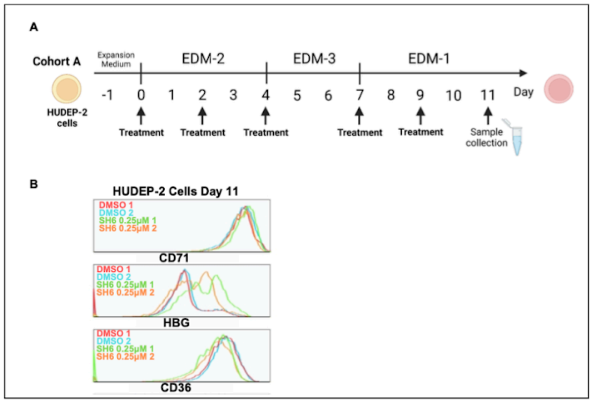


Fig S3. SH6 induces HbF in WT HUDEP-2 cells during a 11 day differentiation protocol. (A) Schematic of HUDEP-2 differentiation and drug treatment following a classic, EDM2 (4 days), EDM3 (3 days) and EDM1 (4 days) differentiation protocol. (B) Flow cytometric histograms of erythroid differentiation markers (CD71, CD36) and HbF in HUDEP-2 cells collected at day 11 of differentiation, following SH6 0.25 μM vs DMSO treatment (n=2 biological replicates each condition).

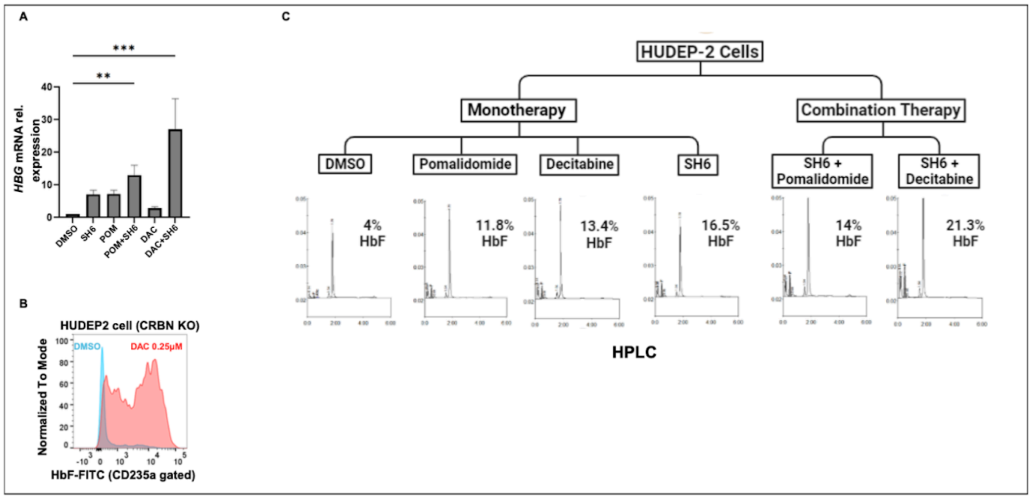

Fig S4. SH6 demonstrates an additive effect of HbF induction with a hypomethylating agent. (A) qPCR analysis of *HBG1/2* mRNA levels following monotherapy treatment with SH6 0.25 μM, Pomalidomide (POM) 1 μM, Decitabine (DAC) 0.25 μM, as well as combination therapy of SH6+DAC, and SH6+POM, in WT HUDEP-2 cells following a 4 days differentiation protocol (n=2 biological replicates, each with 3 technical replicates). (B) Representative flow cytometric analysis of HbF (CD23a+) by DAC in CRBN KO HUDEP-2 cells. (C) Representative HPLC analysis of HbF in WT HUDEP-2 cells treated monotherapies vs combination therapies as described in A.

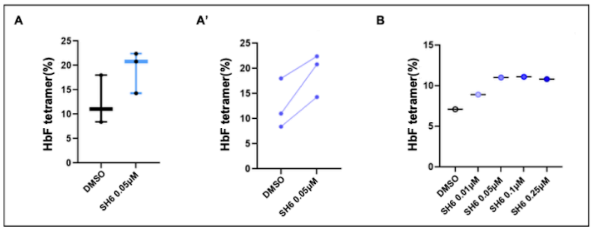

Fig S5. SH6 induces HbF in CD34+ cell-derived erythroid cells. (A-A') Quantification HPLC analysis of HbF in differentiated healthy donor CD34+ cells treated with 0.05 μM SH6 vs DMSO, day 15-17 of differentiation (n= 2 donors, each with 1-2 biological replicates). A' shows paired DMSO and SH6 treatment group, for each biological replicate. (B) HPLC analysis of HbF in terminally differentiated (day 19) SCD CD34+ cells treated with varying doses of SH6 (n=1 patient, 1 biological replicate only due to limited specimen).

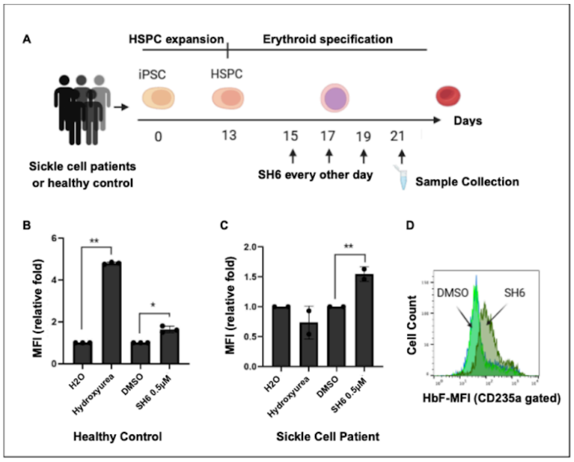


Fig S6. SH6 induces HbF in iPSC-derived erythroid cells from hydroxyurea resistant SCD patient. (A) Schematic of SCD patients and healthy donor iPSC-erythroid differentiation and SH6 treatment, using a 21 day HSPC expansion and erythroid expansion/maturation protocol. (B) Flow cytometric quantification of HbF (relative MFI fold change, CD235a positive cells) of healthy control iPSC-differentiated erythroid cells, treated with hydroxyurea vs. H20 control, or SH6 0.5 μM vs DMSO control (n=2 biological replicates). (C) Flow cytometric quantification of HbF (relative MFI fold change, CD235a positive cells) of SCD iPSC-differentiated erythroid cells, treated with hydroxyurea vs. H20 control, or SH6 0.5 μM vs DMSO control (n=2 biological replicates). The SCD patient was known to be resistant to hydroxyurea. (D) Representative flow cytometric analysis of HbF (CD235a positive cells) of SCD iPSC-differentiated erythroid cells, treated with SH6 0.5 μM vs DMSO control.

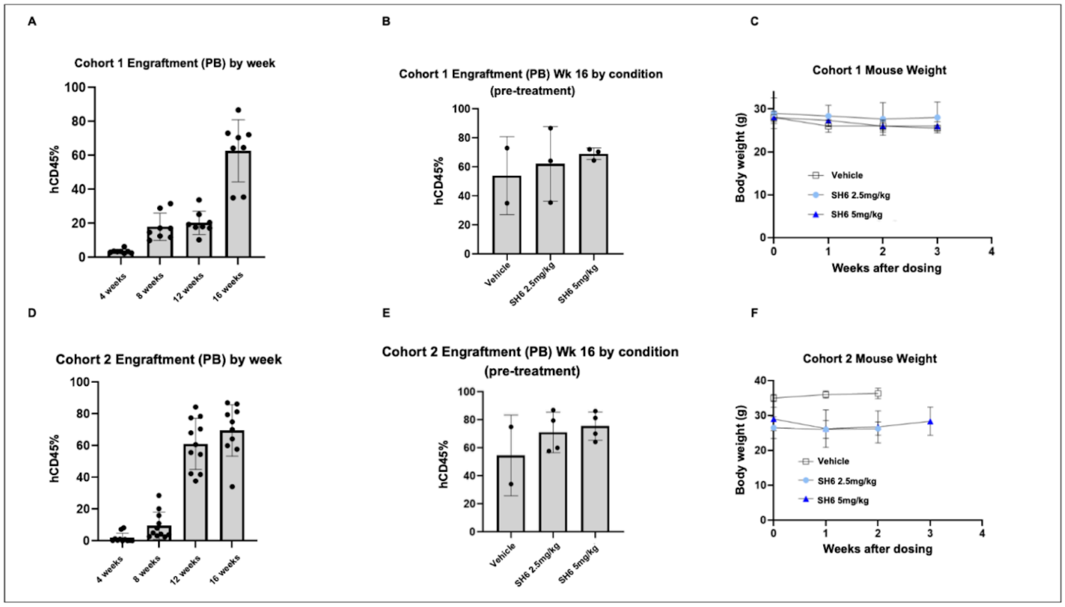

Fig S7. Engraftment trajectory prior to drug treatment and animal weight during drug treatment in the xenotransplantation study. (A) PB engraftment (hCD45%/mCD45+hCD45) in cohort 1 of transplanted mice throughout the engraftment period, assessed every 4 weeks until week 16. (B) PB engraftment (hCD45%/mCD45+hCD45) in cohort 1 of transplanted mice, on week 16 prior to drug treatment, divided by treatment groups (vehicle, 2.5mg/kg, 5mg/kg). (C) Cohort 1 animal weight measurement during the 3 week drug treatment period, across three treatment groups (vehicle, 2.5 mg/kg, 5 mg/kg). (D) PB engraftment (hCD45%/mCD45+hCD45) in cohort 2 of transplanted mice throughout the engraftment period, assessed every 4 weeks until week 16. (E) PB engraftment (hCD45%/mCD45+hCD45) in cohort 2 of transplanted mice, on week 16 prior to drug treatment, divided by treatment groups (vehicle, 2.5 mg/kg, 5 mg/kg). (F) Cohort 2 animal weight measurement during the 3 week drug treatment period, across three treatment groups (vehicle, 2.5 mg/kg, 5 mg/kg).

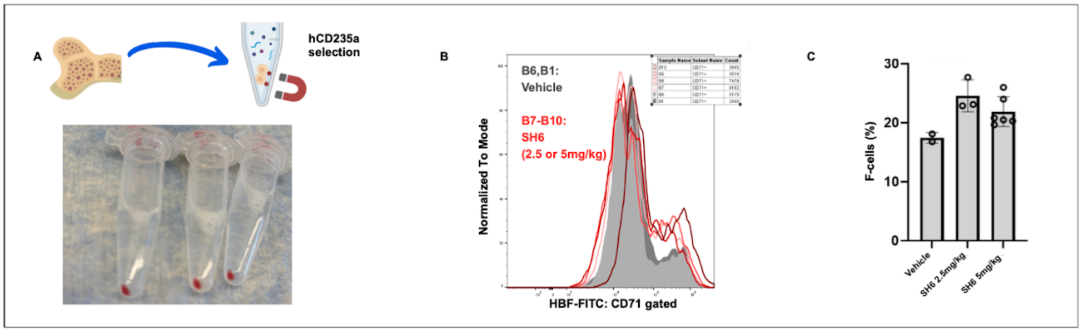


Fig S8. SH6 induces HbF *in vivo* in a human CD34+ xenotransplantation model. (A) hCD235a magnetic bead selection from mouse bone marrow. The pellets represent successfully selected human erythroid cells. (B) Representative flow cytometric analysis of HbF (CD235a and CD71 positive cells) from hCD235a-selected human erythroid cells (grey: vehicle treated; red: SH6 treated). (C) Quantification of flow cytometric analysis of HbF (CD235a and CD71 positive cells) from hCD235a-selected human erythroid cells across three treatment groups (vehicle, SH6 2.5mg/kg, SH6 5mg/kg).

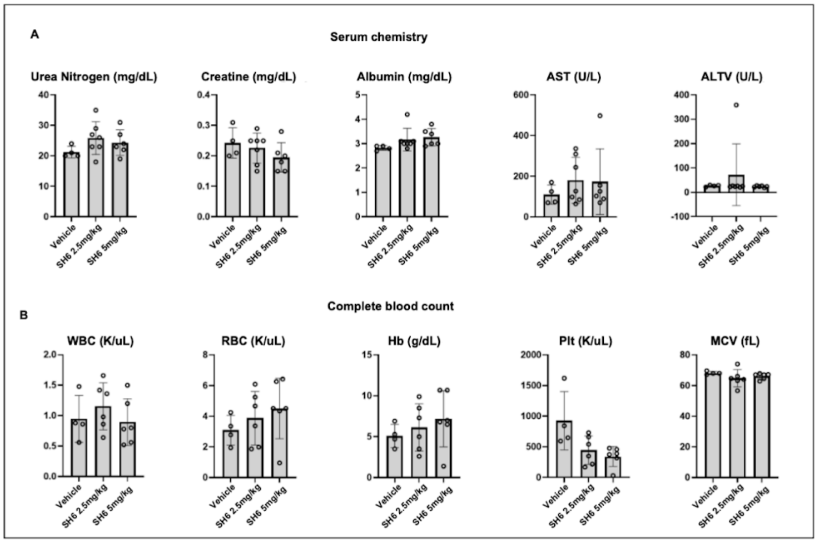


Fig S9. SH6 demonstrates minimal toxicity in a mouse xenotransplantation model. (A) Mouse peripheral blood serum chemistry analysis (Urea Nitrogen, Creatinine, Albumin, AST, ALT) in three treated groups (vehicle, 2.5mg/kg, 5mg/kg) (B) Mouse complete blood count analysis (WBC, RBC, Hb, Plt, MCV) in three treated groups (vehicle, 2.5 mg/kg, 5 mg/kg).

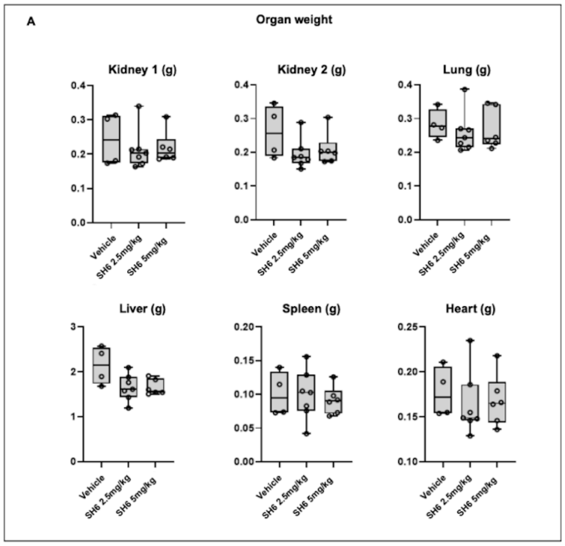


Fig S10. SH6 demonstrates minimal toxicity in a mouse xenotransplantation model. (A) Mouse organ weight (kidney, lung, liver, spleen, heart) in three treated groups (vehicle, 2.5 mg/kg, 5 mg/kg).

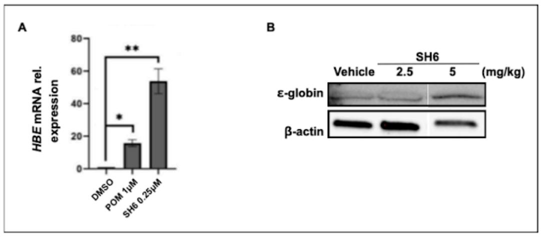


Fig S11. SH6 induces embryonic epsilon globin. (A) qPCR analysis of *HBE* (epsilon globin) mRNA levels following treatment with SH6 and Pomalidomide (POM) vs DMSO in WT HUDEP-2 cells following a 4 days differentiation protocol. (B) WB analysis of epsilon globin protein in hCD235a-selected erythroid cells from mouse bone marrow from healthy donor CD34+ cell xenotransplantation experiment (vehicle, 2.5 mg/kg dose, and 5 mg/kg dose, respectively).

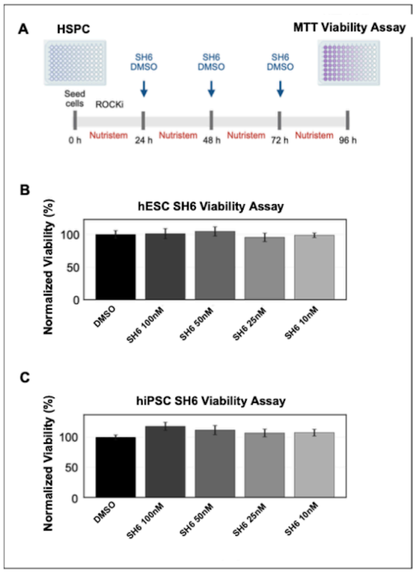


Fig S12. SH6's effects on the viability of human pluripotent stem cells.
(A) Schematic overview of the MTT assay used to measure cell viability after SH6 treatment. (B) Human embryonic stem cells (hESCs) and (C) human induced pluripotent stem cells (hiPSCs) treated with increasing concentrations of SH6 for 72 hours up to 100 nM. Viability data are shown as mean ± SD.

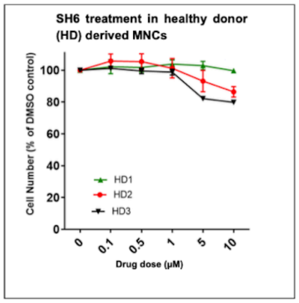


Fig S13. SH6's effects on the viability of healthy donor (HD)-derived mononuclear cells (MNCs). Three individual HD MNCs are treated with escalating doses of SH6 for 24 hours up to 10 μM. Viability data are shown as mean ± SD.

### Supp Table 1. qPCR primer sequences.

HBG1/2 FW TGGCAAGAAGGTGCTGACTTC

HBG1/2 RV GCAAAGGTGCCCTTGAGATC

HBB FW TGGGCAACCCTAAGGTGAAG

HBB RV GTGAGCCAGGCCATCACTAAA

HBE1 FW TGCACTGTGACAAGCTGCAT

HBE1 RV CCTTGCCAAAGTGAGTAGCC

HBA FW AAGACCTACTTCCCGCACTTC

HBA RV GTTGGGCATGTCGTCCAC

GAPDH F ACCACAGTCCATGCCATCACT

GAPDH R CCATCACGCCACAGTTTCC

ZBTB7A F GCAACATCTGCAAGGTCCGCTT

ZBTB7A R TCTTCAGGTCGTAGTTGTGGGC
